# Spontaneous Perceptual Reversals reflect in Internal Rhythms, Not Metabolic Shifts or Evoked Responses

**DOI:** 10.64898/2026.09.09.750323

**Authors:** Lisa Stetza, Lena Hehemann, Christoph Kayser

## Abstract

The perceptual interpretation of a static image, such as the Necker cube, can change spontaneously, resulting in endogenously driven changes in perception. We studied how different peripheral markers of the body’s physiological state, as well as time-resolved markers of EEG-derived brain activity, modulate around such spontaneous changes in perception. Specifically, we quantified the time course of heart rate, respiration rate, respiration phase, pupil size, EEG evoked responses, and EEG time-frequency activity around spontaneous perceptual reversals, and tested how these signals differ between the two interpretations of the Necker cube. Our results show that signals relating to the body’s and brain’s momentary state differ between the two perceptual interpretations, including pupil size, respiration phase, and rhythmic EEG activity. In contrast, signals relating to metabolic state or external stimuli, including respiration rate, heart rate and EEG evoked responses, did not differ between perceptual interpretations, but were consistently modulated around these. This corroborates the notion that spontaneous changes in perception are related to specific and coordinated windows in the cardiorespiratory cycle and neural markers of arousal or top-down processing. These windows are differentially expressed across interacting physiological systems, highlighting the intricate link between perception, brain activity and bodily physiology.

## Introduction

Changes in perception are typically accompanied by changes in brain activity, but also by changes in physiological signals, such as the heartbeat, the respiratory cycle or pupil size. Attempting to understand how specifically these signals relate to perception, many studies specifically test how these signals relate to individual perceptual interpretations or perceptual states (Zelano et al., 2016; Kluger et al., 2021; Grund et al., 2022; Johannknecht & Kayser, 2022; Schaefer et al., 2023; Goheen et al. 2024; Schaefer et al., 2025; Kayser et al., 2026; Skog et al., 2026). Of particular interest in this regard are paradigms that allow dissociating the external stimuli from their perceptual impressions, such as paradigms involving ambiguous stimuli (Klink et al., 2012; Devia et al., 2022). For example, when viewing a static image of a Necker cube our perceptual interpretation of this changes back and forth between the two interpretations of the cube. These so-called perceptual reversals are accompanied by systematic changes in brain activity (Britz et al., 2009; Sterzer et al., 2009; Knapen et al., 2011; Kornmeier & Bach, 2012; Wang et al., 2013; Kloosterman et al., 2015; Brascamp et al., 2018; de Jong et al., 2020; Wilson et al., 2023) and pupil size (Einhäuser et al., 2008; Hupé et al., 2009; de Hollander et al., 2018; Sato et al., 2020; Brascamp et al., 2021; Nakano et al., 2021). While brain activity has been frequently studied in this context (Devia et al., 2022), much less is known about changes in bodily physiology accompanying perceptual reversals. We here address this gap by providing a comprehensive analysis of human EEG-derived neural and peripheral physiological signals (respiration, ECG, pupil) around spontaneous changes in perception.

One line of previous studies has tested how neurophysiological signals, e.g. measured using EEG or fMRI, relate to changes in the conscious interpretation of sensory stimuli. This has shown that perceptual reversals are characterized by slower and more gradual changes in neural activity than changes in percepts induced by suddenly presented stimuli (Sterzer et al., 2009; Wang et al., 2013; Brascamp et al., 2018; Devia et al., 2022). While neural activity related to endogenously driven changes in percepts often involves frontal, temporal, and parietal regions, some studies also implied early sensory regions (Pitts et al., 2010; Sanders et al., 2018; Curtu et al., 2019). Furthermore, studies of rhythmic brain activity have reported a decrease in alpha-band power over occipital-parietal regions starting about one second prior to the reversal, possibly associated with reduced inhibitory processes facilitating subsequent changes in perception (İşoğlu-Alkaç & Strüber, 2006; Piantoni et al., 2010; Piantoni et al., 2017; Drew et al., 2022; Mokri et al., 2025; Drew et al., 2026). Similarly, beta-band power was shown to decrease prior to (Piantoni et al., 2010; Piantoni et al., 2017) and increase subsequent to a reversal (Piantoni et al., 2010; Yokota et al., 2014; Piantoni et al., 2017) and changes in theta-band activity were associated with the resolution of perceptual conflicts (Drew et al., 2022; Drew et al., 2026). Importantly, for the Necker cube, the two perceptual interpretations of the ambiguous drawing are not fully equivalent. One interpretation is considered more ‘natural’, as it resembles the objects’ orientation more frequently experienced in real life and this perceptual interpretation typically remains stable for longer periods than the alternative interpretation (Nakayama & Shimojo, 1992; Kornmeier et al., 2009; Sato et al., 2020). Some studies show that based on neurophysiological signals it is possible to differentiate between these two perceptual interpretations, suggesting that the dynamics of neural signals allows reading out the individual perceptual state (Hramov et al., 2017; Gelbard-Sagiv et al., 2018; Devia et al., 2022; Pitts et al., 2010; Sanders et al., 2018; Curtu et al., 2019).

Another line of studies has shown that endogenously driven changes in perception are accompanied by changes in pupil size, a peripheral signal that reflects arousal, attention and cognitive effort (Aston-Jones & Cohen, 2005; Laeng et al., 2012; Viglione et al., 2023) and is related to activity of the locus coeruleus (Aston-Jones & Cohen, 2005). Previous work shows that pupil size systematically constricts prior to a perceptual reversal and dilates subsequently, suggesting that multiple and interacting processes underlie changes in perception (Einhäuser et al.,2008; Hupé et al., 2009; Brascamp et al., 2021; Nakano et al., 2021). As for neural activity, some studies show that the precise dynamics of the pupil size differs between the two perceptual interpretations, suggesting that the physiological state may depend or change with the current percept (Sato et al., 2020).

In contrast, less is known about how cardiac or respiratory activity change around perceptual reversals. Many studies have described that respiration phase aligns to the repeated presentation of external stimuli, resulting in a specific respiratory phase to prevail around stimulus onset (Huijbers et al., 2014; Perl et al., 2019; Kluger et al., 2021; Johannknecht & Kayser, 2022; Goheen et al., 2024; Andrews et al., 2025; Stetza et al, 2025; Chalas et al., 2026; Harting et al., 2026; Kayser et al., 2026; Saltafossi et al., 2025), while only few accounts investigated possible effects of other respiratory parameters as well (Kosik-Rose et al., 2026; Skog et al., 2026). This alignment of respiration to external inputs could reflect the preparatory modulation of cortical excitability to optimize neural processing of expected stimuli (Kluger & Gross, 2021; Della Penna et al., 2026; Chalas et al., 2026). In contrast to externally-paced stimuli, little is known about how respiration relates to endogenous changes in perception. Two studies using binocular rivalry showed that the probability for a reversal to occur is highest during expiration and that changes in respiration rate can affect the precise time of highest reversal probability (Yonekura et al., 2025; Iwamoto et al., 2025). In a previous study, we showed respiration phase-locking occurring after the reversal, although relatively weak when compared to phase-locking to external stimuli (Stetza & Kayser, 2026).

The same also applies to cardiac activity. Anticipatory cardiac deceleration precedes expected stimuli (Skora et al., 2022; Kingir et al., 2026; Jennings et al., 1977; Jennings & Van der Molen, 2005; Motyka et al., 2019) and behavioral performance depends on the cardiac phase (systole vs. diastole), affecting the detection of near-threshold stimuli (Grund et al., 2022; Motyka et al., 2019; Al et al., 2020; Al et al., 2021). Similarly, self-initiated motor action (Kunzendorf et al., 2018, Gerosa et al., 2026) and cortical motor excitability (Al et al., 2023) were shown to vary with cardiac phase. Similar to respiration, cardiac activity has been little studied in the context of spontaneous perceptual reversals. One study showed that perceptual dominance during binocular rivalry is influenced by the cardiac phase (Veillette et al., 2024), while another study reported a dependency between perceptual uncertainty and heart rate in binocular rivalry (Corcoran et al. 2021). Heart-beat related neural processes can also be measured using the heartbeat-evoked potential (HEP), which reflects the cortical processing of cardiac afferent signals and is often used as electrophysiological marker of interoceptive processing (Schandry et al., 1986; Schandry & Weitkunat, 1990; Pollatos & Schandry, 2004). While HEPs have been related to the processing of sensory stimuli (Park et al., 2014; Azzalini et al., 2019; Al et al., 2020; Al et al., 2021), and cortical motor excitability (Al et al., 2023), little is known about their relation to endogenously driven changes in perception.

In sum, spontaneous changes in perception provide an important testbed to understand the neural and bodily physiology underlying conscious perception. Importantly, by avoiding exogenously induced changes in the physical input, paradigms such as the Necker cube allow linking physiological states to endogenously induced changes in perceptual interpretations. While several studies have shown that neural and bodily physiological signals are modulated around perceptual reversals, most studies focused on a single signal, precluding a direct comparison between the insights obtained from different measurements of neural and bodily states. Hence, we performed a comprehensive analysis of how respiration phase, respiration frequency, heart rate, pupil size, EEG-derived event-related potentials, EEG time-frequency dynamics, and heartbeat-evoked potentials change around perceptual reversals of the Necker cube. We specifically asked whether and which of those signals differ between the two perceptual interpretations of the cube, and hence possibly allow differentiating the perceptual state of the participant.

## Methods

### Participants

The study included 35 adult volunteers who provided written informed consent prior to participation. All participants reported having normal or corrected-to-normal vision and hearing and received compensation for their participation. The sample mainly consisted of young university students with typical demographics, although no demographic information was collected as the data was collected anonymously. Participants were briefed on the experimental setup and equipment; however, they were not specifically informed that the investigation examined the association between respiration, cardiac activity and pupillometry during bistable perception, in line with our earlier studies (Johannknecht & Kayser, 2022; Stetza et al., 2025; Kayser et al., 2026; Stetza & Kayser, 2026). They were asked to breathe normally through their nose, although oral breathing during some parts of the experiment cannot be ruled out. The study protocol was approved by the Ethics Committee of Bielefeld University.

### Experimental setup and data acquisition

The experiment was performed in a sound-attenuated and electrically shielded booth (Desone, Germany). Visual stimuli were displayed on a 27-inch monitor (ASUS PG279Q, 120 Hz) placed at an approximate viewing distance of 85 cm from the participant. Stimulus presentation was controlled with MATLAB (R2017a, MathWorks Inc., Natick, MA) in combination with Psychtoolbox (Version 3.0.14, Kleiner et al., 2007). No additional light sources were present during the experiment apart from the monitor. Synchronization between visual stimulus presentation and physiological recordings was achieved through TTL pulses transmitted to the BioSemi ActiveTwo system (ActiView software).

Respiratory data were measured using a temperature-sensitive thermistor (Littelfuse GT102B1K, Mouser Electronics) mounted on a modified disposable oxygen mask (Johannknecht & Kayser, 2022; Harting et al., 2026; Stetza & Kayser, 2026). Voltage changes caused by temperature variations associated with nasal airflow during inhalation and exhalation were amplified and acquired through the analogue input channels of the BioSemi EEG system. Eye movements and pupil diameter were recorded from the left eye using an EyeLink 1000 system (SR Research Ltd., Canada) at 500 Hz in free-viewing mode. Before each experimental block, a 9-point calibration procedure was carried out. Pupil diameter was determined using the system’s Centroid algorithm. The ECG was recorded using the ActiveTwo system using two Ag-AgCl surface electrodes attached just below the bend of the arm, with one electrode attached to each arm. The CMS and DRL electrodes on the EEG cap served as reference and ground for the ECG. EEG data were acquired with a 128-channel system at a sampling frequency of 1024 Hz, using Ag-AgCl electrodes embedded in an elastic cap (BioSemi BV). Electrode offsets were maintained within a range of ±25 mV. Reference electrodes were positioned at their standard locations on both sides of the midline, slightly posterior to the Cz electrode. Electrooculographic (EOG) activity was recorded using four Ag-AgCl surface electrodes placed laterally and below each eye.

### Experimental task

The experimental paradigm and stimuli were identical to our previous study (Stetza & Kayser, 2026). A Necker cube (11° × 9° of visual angle; line width: 3 pixels; RGB 115, 115, 115) was presented on a uniform grey background (RGB 70, 70, 70; 18.5 cd/m²) and remained constant during both stimulus presentation and inter-trial intervals to avoid luminance changes. The paradigm structure consisted of 36 trials of 75 seconds each, with 15-second inter-trial intervals. Trials were organized into six blocks, each containing six trials, with breaks between blocks. Prior to each block, participants were adapted to darkness to minimize pupil size fluctuations related to changes in ambient light.

Participants were instructed to maintain their gaze on the screen to ensure they did not miss stimulus onset or offset but were otherwise allowed free viewing to reduce fatigue associated with sustained fixation. They were not given instructions regarding specific gaze strategies or intentional modulation of perceptual reversals and were kept naïve to the study purpose as far as possible. The task instruction was to press a key on the computer keyboard whenever the perception of the cube changed, with the left and right arrow keys associated with the two possible interpretations of the cube (Stetza & Kayser, 2026). These two interpretations were illustrated and practiced by the participants in a block prior to the actual experiment.

### Signal Preprocessing

#### EEG

EEG data were preprocessed using FieldTrip (Oostenveld et al., 2011) in MATLAB (R2022b). Signals were band-pass filtered using a 0.6 Hz high-pass (4th order) and a 90 Hz low-pass filter, followed by resampling to 200 Hz. Bad channels (1.5 ± 1.5, mean ± SD) were identified and interpolated, and standard preprocessing routines were applied in FieldTrip (*ft_preprocessing, ft_resample, ft_prepare_neighbours, ft_channelrepair*). Subsequently, independent component analysis (ICA) was performed using the ‘runica’ algorithm (*ft_componentanalysis*) after performing PCA with 40 principal components. Artifact-related components were removed based on correlations with movement artifacts, topographical features, contributions of EOG and ECG channels, as well as spectral characteristics indicative of low LF/HF ratios as described previously (Kayser et al., 2017; Grabot & Kayser, 2020) (11.9 ± 4.5 removed components, mean ± SD). Data were then segmented into epochs time-locked to perceptual reversals, with additional exclusion of epochs exceeding ±200 µV.

#### ECG

ECG signals were filtered with a 2 Hz high-pass filter, a 40 Hz low-pass filter, and a 49–51 Hz band-stop filter, followed by resampling to 200 Hz. Movement-related artifacts were reduced using *findpeaks* with a minimum peak height defined as mean(signal) + 0.5 × SD (signal) and a minimum peak distance corresponding to 0.4 times the resampling frequency. R-peaks were subsequently identified using a Pan-Tompkins-based algorithm using the *ft_heartrate* function in FieldTrip. An additional quality check ensured that only heartbeats with inter-beat intervals greater than 650 ms were included in the analysis. Again, the data were then segmented into epochs time-locked to perceptual reversals.

### Respiration

Respiration data were preprocessed in MATLAB (R2022b) similar to our previous work (Johannknecht & Kayser, 2022; Harting et al., 2026; Stetza & Kayser, 2026). Signals were filtered using a third-order Butterworth filter (0.03 Hz high-pass, 6 Hz low-pass), resampled to 100 Hz, and converted to z-scores. Respiration cycles were defined around each peak (Johannknecht & Kayser, 2022; Harting et al., 2026; Stetza & Kayser, 2026). For some respiratory cycles, brief pauses following exhalation were classified as a third respiratory state and were therefore excluded from further analyses (Noto et al., 2018). For analysis, respiration phase was defined as a circular-linear variable progressing from 0 to π during inspiration and from π to 2π during expiration, providing a continuous representation with meaningful reference points (0/2π = peak inhalation; π = peak exhalation). Besides respiratory phase, we also stored the moment-by-moment duration of respiration cycles as a continuous trace reflecting respiration rate.

### Pupil Size

The pupil data were processed as follows. First, the eye tracking data were cleaned by removing outliers in gaze coordinates (>14 degrees) as well as samples exceeding thresholds in eye movement acceleration (> 800 degree/s) or velocity (>1.2 degree/s^2^), followed by z-scoring. Blinks and missing data segments shorter than 100 ms were linearly interpolated. Slow temporal drifts were corrected by fitting and subtracting an exponential function, and the signal was low-pass filtered at 8 Hz and resampled at 100 Hz to reduce high-frequency noise, as suggested previously (Brascamp et al., 2021; Stetza & Kayser, 2026).

### Data exclusion and cleaning

Based on the behavioral data, one participant had to be excluded due to a very low number of reported perceptual reversals (N = 34). Similar to our previous work, we excluded consecutive presses of the same response button, epochs with more than 25 s between adjacent reversals, and trials that contained only a single epoch.

We analyzed each of the different signals independently in order to maximize the number of available participants and data epochs around perceptual reversals for each analysis. Hence, for each signal, we counted the number of available data epochs and excluded participants who had too few epochs for this specific signal, using a cut-off of 120 reversals.

For respiration, two participants were excluded due to faulty data recordings, and one was excluded based on the number of available remaining data epochs. Hence, we report data for N = 31 participants (497.8 ± 281.9 epochs, mean ± SD). Overall, 74.3 ± 16.3 % of all reported reversals were retained in this analysis. For eye tracking, one participant had to be excluded based on the number of available data epochs, and we report data from N = 33 participants (591.3 ± 280.4 epochs, mean ± SD). Overall, 91.3 ± 6.1 % of the reported reversals were retained in the analysis. For the EEG data, we retained data from N = 29 participants, with 270.3 ± 90.9 epochs (mean ± SD), which correspond to 55.9 ± 19.0 % of all reported reversals. For the ECG data, data from 7 participants had to be excluded as an electrode lost contact, rendering the R peak detection unreliable or impossible. We report data from N = 27 participants (699.2 ± 307.5 epochs).

For each of the signals, we report the average time course of this signal around the perceptual reversal. We also tested whether the signal differs between the two perceptual interpretations of the cube. In line with previous observations that the two interpretations of the cube differ in their stability and prevalence, we label the right-oriented interpretation as non-dominant (yellow in Figure 2A) and the left-oriented as dominant (blue in Figure 2A). In line with this, we label the perceptual reversals in which the interpretation changes from right-oriented to left-oriented cube as ‘*to dominant*’, and those from left- to right-oriented as *‘to non-dominant*’.

**Figure 1:**
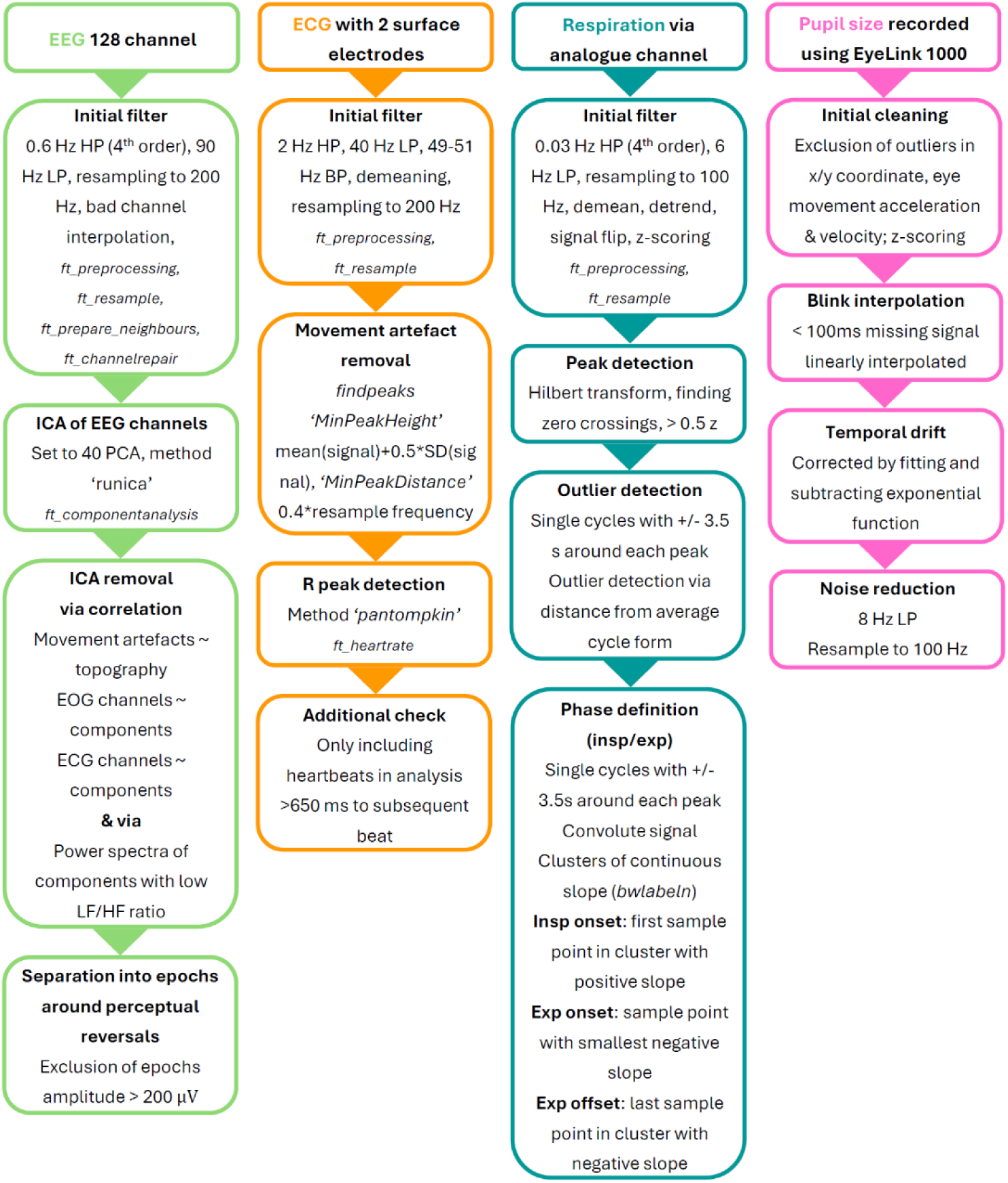
Flow chart of preprocessing steps for all four signals. The chart lists the key steps, including relevant functions in Matlab (in italics) and thresholds for filtering or data exclusion.

**Figure 2:**
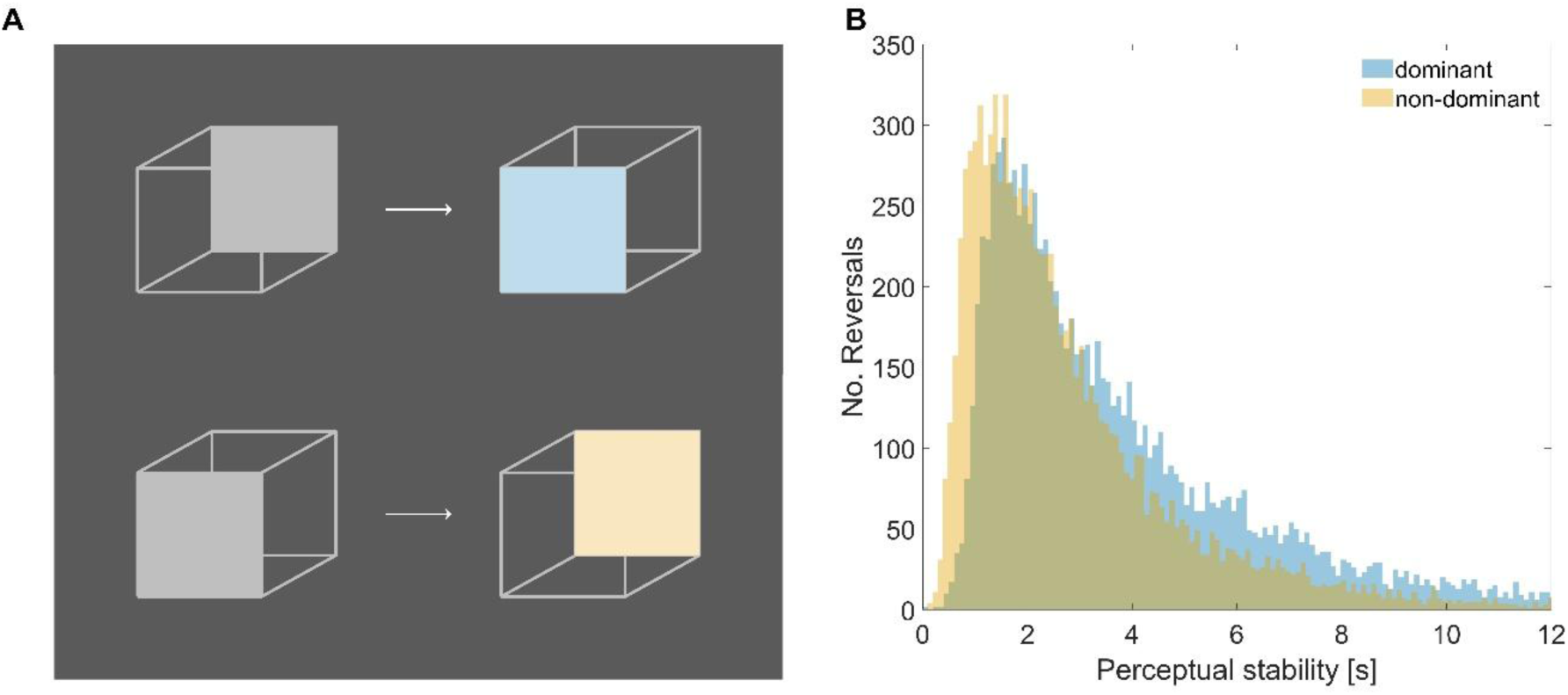
**A)** Example of the two perceptual reversals. In the upper example the percept changes from the non-dominant (left-oriented) interpretation to the dominant interpretation (‘to dominant’; blue); in the lower example it changes to the non-dominant interpretation (‘to non-dominant’; yellow). **B)** Distribution of the perceptual stability prior to each reversal across all participants and reversals. The dominant interpretation was consistently seen longer, with a perceptual stability of 4.10 ± 3.35 s (mean ±SD) for the dominant cube, and 2.92 ± 2.41 s (mean ±SD) for the non-dominant cube.

### Analysis of respiration data and ECG

To analyze respiration phase, we followed our previous work and quantified the alignment of respiration phase to the reversals using the phase-locking value (PLV) (Johannknecht & Kayser, 2022; Harting et al., 2026; Stetza et al., 2025; Kayser et al., 2026; Stetza & Kayser, 2026). For this, the respiration phase was transformed into complex numbers and averaged across epochs within participants. The PLV is then obtained as the vector length of this average. To test whether the group-average of the PLVs differs from zero, we relied on permutation testing (Park et al., 2020; Johannknecht & Kayser, 2022). For this, we obtained a distribution of group-averages of the PLV under the null hypothesis of no temporal alignment of respiration to the reversal. Practically, we randomly shifted the respiration data relative to the reversal and obtained a surrogate distribution of group-averages generated from 4000 random temporal shifts implemented independently for each participant (Johannknecht & Kayser, 2022; Harting et al., 2026; Stetza et al., 2025; Kayser et al., 2026; Stetza & Kayser, 2026; Kluger & Gross, 2021). To correct for multiple comparisons along time, we extracted the maximal group-average PLV across all time points for each surrogate sample. We then computed the p-value of the actual PLV based on the percentile of the surrogate distribution.

To analyze respiration rate, we extracted the momentary duration of respiration cycles and averaged these across reversals within participants. To visualize the time course of cycle duration around reversals, we z-scored the epoch-averaged data within each participant to normalize between-participant differences in respiration rate (Figure 3F). To compare the duration of respiration cycles between reversals, we contrasted the epoch-averaged signal, without z-scoring. To analyze heart-rate, we calculated the epoch-averaged rate for each participant and proceeded similarly as for the cycle duration, relying on a z-scored signal to visualize the overall time course around the reversal. Finally, for pupil size we calculated the epoch-averaged signal, both across all reversal and for each reversal separately, without any further standardization.

**Figure 3:**
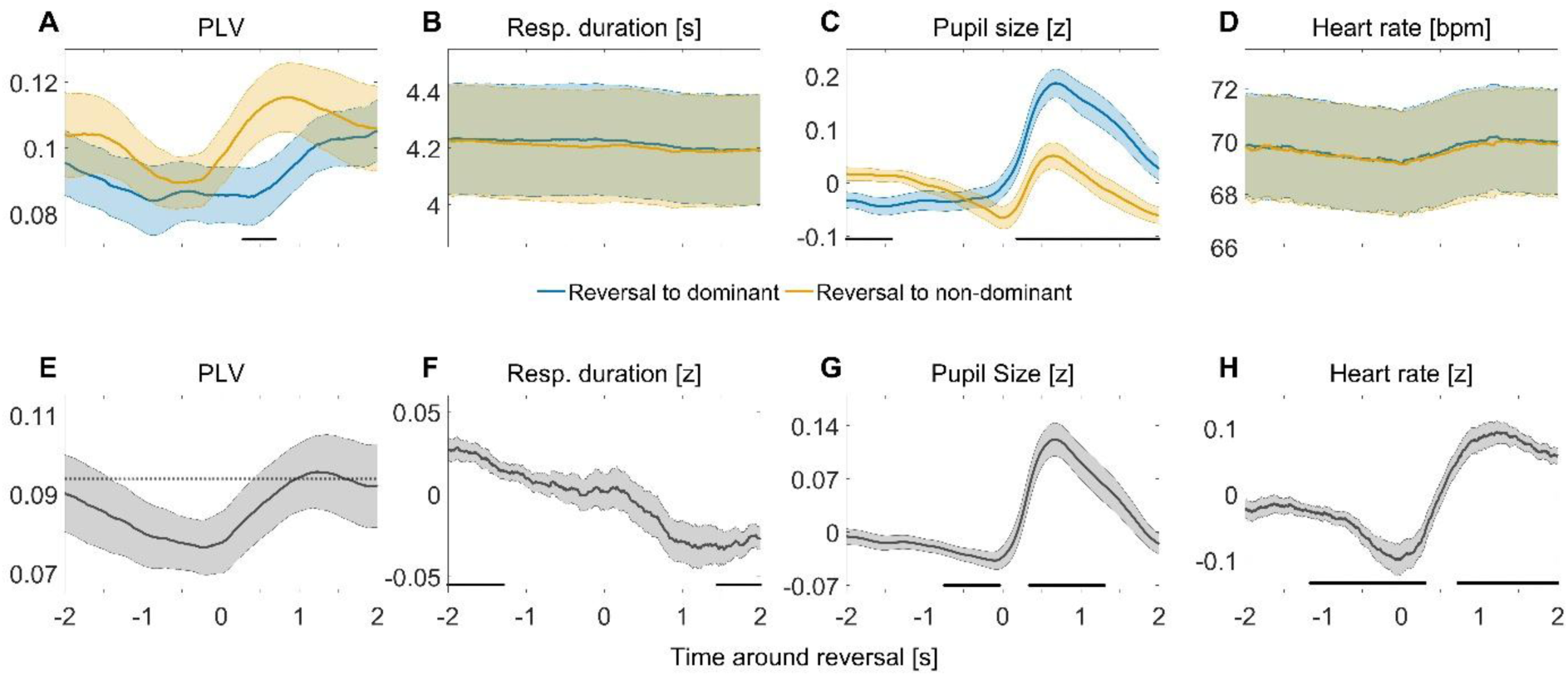
Peripheral signals around perceptual reversals of the Necker cube. **A)** Phase locking of respiration phase relative to reversals (at t=0). The yellow trace shows the reversal to the non-dominant interpretation (max PLV = 0.115 at 0.88 s), the blue trace the to-dominant reversal (max PLV = 0.105 at 2.0 s). Thick lines indicate the group-average, shaded areas the standard error. The PLV differs significantly between reversals between 0.28 to 0.69 s after reversal (cluster-based permutation test; t = -3.0, p = 0.0025). **B)** Duration of respiration cycles (i.e. inverse respiration frequency). This did not differ between reversals. **C)** Pupil size. This differed significantly between reversals (cluster 1: -2 to -1.42 s; t =-3.0706, p = 0.0025; and cluster 2 from 0.18 to 2.0 s, t = 5.3935, p = 0.0025). **D)** Heart rate. This did not differ between reversals. **E)** Collapsed PLV across both reversals. PLV reaches significance (p < 0.01; dashed line) between 0.97 to 1.57 s with its maximum of 0.096 at 1.22 s. **F)** Collapsed and normalized respiration cycle duration. This differed significantly from zero (from -2.0 to -1.29 s; t = 4.5632, p = 0.0025, and 1.41 to 2.0 s; t = -4.2142, p = 0.0025). **G)** Collapsed and normalized pupil size across both percepts with significant difference from zero in two clusters, from -0.75 to -0.04 s (t = -3.4634, p = 0.0025) and 0.34 to 1.3 s (t = 5.6901, p = 0.0025). **H)** Collapsed and normalized heart rate (bpm). This differed significantly from zero (from -1.17 to 0.31 s; t = -4.9420, p = 0.0012; and 0.72 to 2.0 s, t = 6.8030, p = 0.0012).

For each of these signals, we tested for statistically significant differences between the two reversal types, based on a cluster-based permutation test correcting for multiple comparisons along time. The first-level contrast was based on paired t-tests across participants, and a surrogate was obtained using 5,000 randomizations of the condition difference (thresholding the first-level *t*-test at α = 0.01; requiring a minimum cluster size of 6; using the max sum as cluster forming statistics).

### Calculation and analysis of event-related potentials

To reduce the influence of confounding motor action of neighboring reversals, we excluded all epochs that had another reversal within a ± 2 s window. This reduced the dataset to N = 29 participants (279.5 ± 91.2 epochs). We then computed the participant-wise epoch-averaged evoked potential, once across all reversals and separately for each type of reversal (using ft_timelockanalysis; Figure 4).

**Figure 4:**
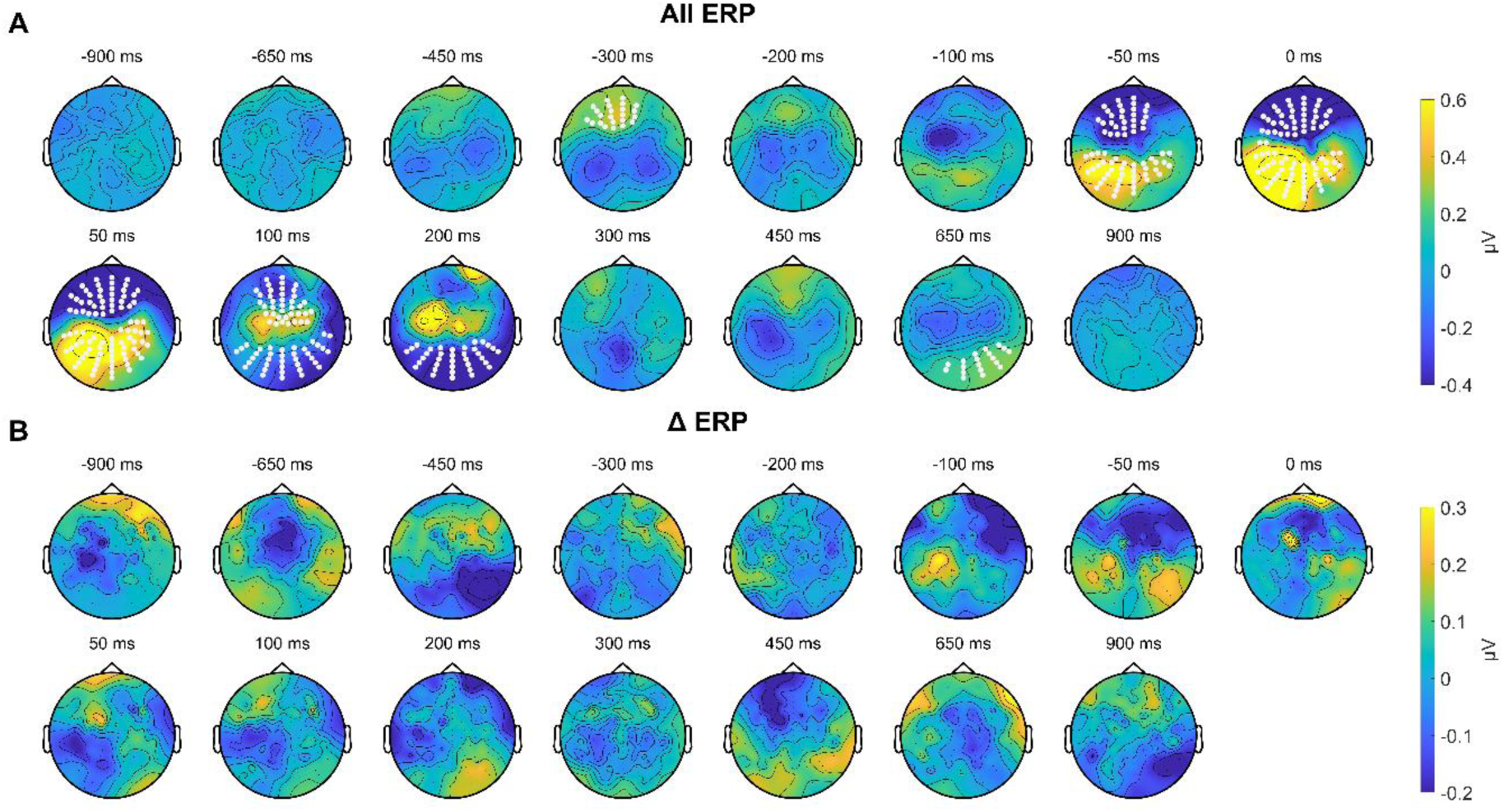
**A)** Average evoked responses across all reversals. The evoked responses differ significantly from zero in several time-electrode clusters indicated by the white dots (see text for details). **B)** Difference in ERPs between reversals. Cluster permutation revealed no significant differences.

To test for differences in ERP between reversals, we used a cluster-permutation scheme relying on paired t-tests across participants and correcting for multiple comparisons along time and electrodes (5000 randomizations, thresholding the first-level *t*-test at α = 0.01; requiring a minimum cluster size of 4 electrode/time points; using the max sum as cluster-forming statistics).

### Time-frequency analysis

As for the ERP analysis, we relied on epochs without another reversal in a window of ± 2 s. To extract rhythmic EEG signals we implemented a time-frequency analysis using the same epochs as used for the ERP (ft_freqanalysis; method = mtmconvol, taper = hanning, output = pow, center frequencies = [4:1:30,32:2:40] Hz, Hanning window of 4.5 cycles). We then log-transformed the power and averaged this across reversals. To quantify differences in the TF-spectrum between conditions, we analyzed the power in five regions of interest (ROI; Figure 5). To identify statistically relevant differences between conditions, we performed a cluster-permutation test with the same parameters as in the ERP analysis.

**Figure 5:**
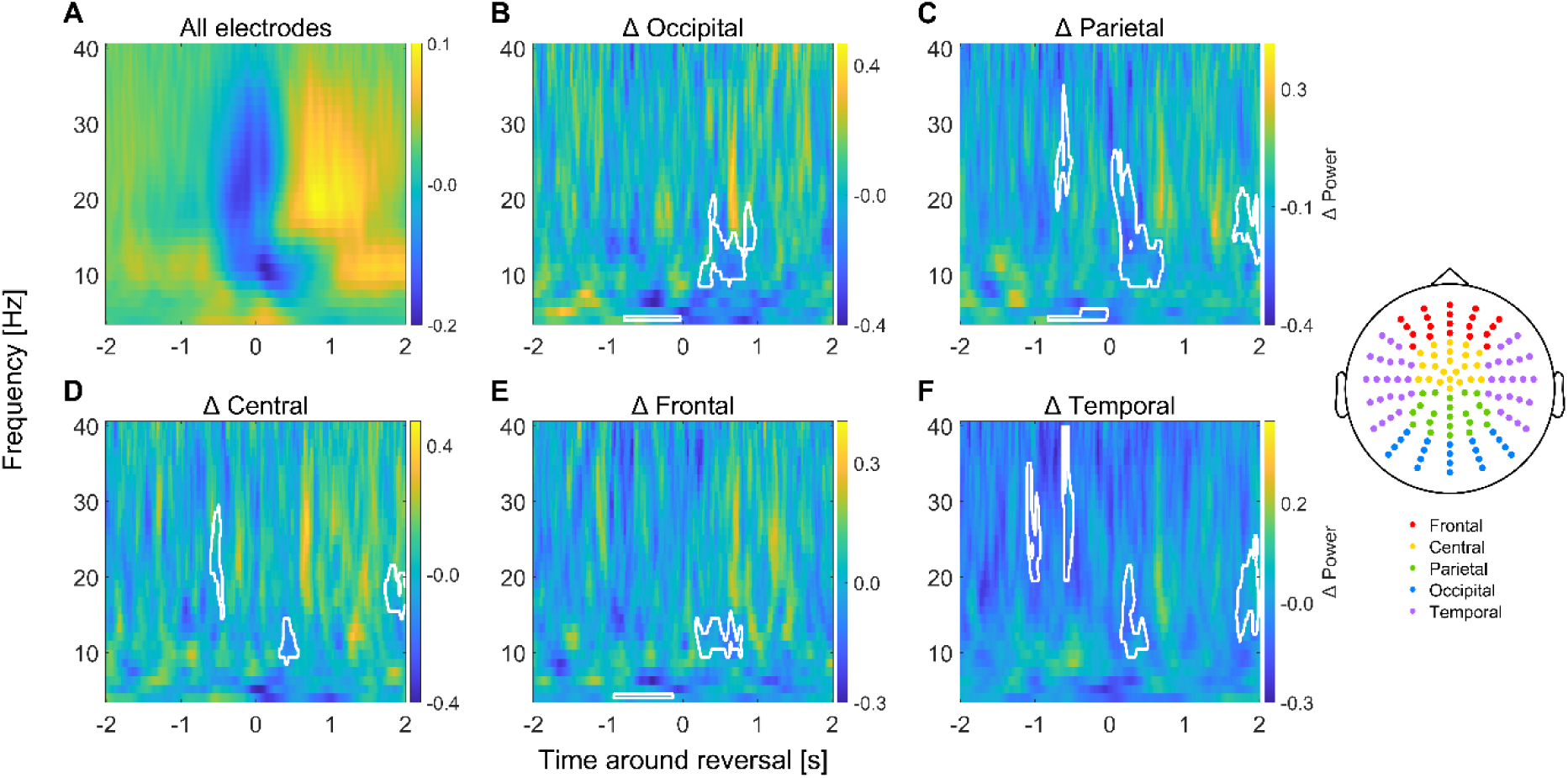
**A)** EEG time-frequency power around reversals, averaged across all electrodes and both types of reversals. To visualize the changes in power, the power spectrum was normalized relative to the average power in the ± 2 s window, separately for each frequency. **B-F)** Difference (to-dominant minus to-non-dominant) in power in five regions of interest. White frames indicate clusters of significant differences between reversals. The topography on the right depicts the electrode positions of the five ROI.

### Heartbeat evoked potential (HEP)

To calculate the HEP, we relied on the R peaks derived from the ECG data and epoched the EEG data around these. For this, we required both valid ECG and EEG data, and this analysis was based on N = 23 participants (deriving 2818.2 ± 476.3 HEPs, mean ± SD). To derive the HEP, we baseline corrected the EEG signal in a window beginning with the R peak until 650 ms after the R peak with the signal’s average in the window of -200:0 ms relative to the R peak (Virjee et al., 2026; Steinfath et al., 2026). To test whether the HEP differs between the two types of perceptual reversals, we detected the last full HEP time window preceding a reversal (allowing a maximum time span of -1.2 prior to the reversal) and the first full HEP window following this. This resulted in an average of 251.3 ± 95.1 reversals (mean ± SD) that could be included in the analysis. To quantify statistically relevant differences between conditions, we applied the same cluster-permutation logic as for the ERP analysis.

## Results

### Respiration alignment but not rate differs between percepts

The consistency of the respiration phase across reversals within each participant was measured using a phase locking index (PLV). This phase consistency was modulated around perceptual reversals and was smaller prior to a reversal and increased subsequently (maximal PLV = 0.096 at 1.24 s). This PLV was statistically significant (compared to a baseline of no significant phase locking) only after the reversal (reaching p<0.01; c.f. Fig. 3F). Interestingly, respiratory phase locking also differed between the two reversals (significant cluster from 0.28 to 0.69 s after the reversal, t = -3.0, p = 0.0025; Fig 3 A) and respiratory phase locking was stronger for the to-non-dominant reversal. We also extracted the individual reversal-averaged respiratory phase angle value at the time of maximum PLV. For reversals to the non-dominant cube, the mean respiration phase was at mid expiration (279.63°) at the time of maximum PLV (at 0.88 s). At the same time point, the average respiration phase during the to-dominant reversals was in late expiration (301.04°).

The average respiration cycle duration in the present data across the entire time was 4.06 ± 1 s (mean ± SD). Around reversals it was slightly higher, reflecting a slight decrease in respiration rate around the reversals. The duration of respiratory cycles also systematically changed around reversals (Fig. 3F). Specifically, the duration of respiratory cycles decreased subsequently to a reversal (avg. prior to reversal 4.23 s/cycle; subsequent 4.19 s/cycle), reflecting an increase in respiratory rate by 40 ms (0.0023 Hz). Statistical testing revealed that the normalized cycle duration differed significantly from zero at multiple time points (cluster from -2.0 to -1.29 s, t = 4.5632, p = 0.0025 and one cluster from 1.41 to 2.0 s; t = -4.2142, p = 0.0025). Contrasting the respiratory cycle duration between the two reversals, however, resulted in no significant differences (at p<0.05; BF_10_ = 1.1 at maximum difference at -0.13 s; Figure 3B).

### Pupil size systematically differs between reversals

Pupil size systematically decreased prior to an indicated perceptual reversal and increased subsequently (Figure 3G), resembling a pattern reported in several studies (Einhäuser et al., 2008; Hupé et al., 2009; de Hollander et al., 2018; Brascamp et al., 2021; Nakano et al., 2021). The normalized pupil signal differed from baseline both prior and subsequent to a reversal (cluster from -0.75 to -0.04 s; t = -3.4634, p = 0.0025; 0.34 to 1.3 s; t = 5.6901, p = 0.0025), corroborating a considerable modulation of pupil size around reversals. Pupil size also systematically differed between the two reversals, in particular subsequent to the reported change (cluster from 0.18 to 2.0 s; t = 5.3935, p = 0.0025): the change to the dominant interpretation was characterized by a more pronounced pupil dilation (Figure 3C, blue). Pupil size also differed significantly prior to the reversal (cluster from -2 to -1.42 s; t =-3.0706, p = 0.0025), where the momentary percept of the dominant interpretation (in the to-non-dominant reversal) was associated with a larger pupil size. All in all, this suggests that the perception of the dominant interpretation of the cube is systematically associated with a large pupil dilation, and changing towards this interpretation also comes with a larger modulation of pupil size.

### Heart rate changes around but does not differ between reversals

The normalized heart rate reveals a deceleration prior to the reversal followed by an acceleration afterwards (Figure 3H). Supporting this, the normalized heart rate differed significantly from zero in two epochs: one corresponding to the deceleration (cluster from - 1.17 s to 0.31 s; t = -4.9420, p = 0.0012) and one corresponding to the acceleration (cluster from 0.72 to 2.0, t = 6.8030, p = 0.0012). Similar to respiration frequency also heart rate did not differ significantly between reversals (at p<0.05; BF_10_ = 1.13 at maximum difference at 1.17 s; Fig. 3D). Heart rate increased from its minimum of 69.14 bpm at -0.01 s to its maximum at 1.22 s of 70.09 bmp which equals a change in heartbeat duration of 11.8 ms.

### Event-related potentials do not differ between reversals

Figure 4A shows the EEG-derived event-related potentials in a window ± 900 ms relative to the reversal. This reveals a change in this neurophysiological signal both prior and subsequent to the reversal, reflecting a specific dynamics of brain activity associated with the change in perceptual interpretation and or the associated motor response. This included a frontal positivity around -300 ms prior to the reversal report (cluster-based permutation test; left-sided fronto-temporal positive clusters -375 to -360 ms, p = 0.0244; -325 to -290 ms, p = 0.0094) and a fronto-occipital difference just around the reversal (occipito-temporal positive cluster -70 to +60 ms, p = 0.0002; frontal negative cluster -55 to +75 ms, p = 0.0002). Around 100 to 200 ms after the reversal report the posterior activity becomes negative (occipito-temporal negative cluster 115 to 195 ms, p = 0.0002) while centro-parietal channels remain positive (fronto-central positive cluster 125 to 160 ms, p = 0.0012), and the frontal negativity subsides. A short positive cluster appears over fronto-central channels 360 to 375 ms after the reversal (p = 0.0342). Late after the reversal, at 650 ms, a positive activation over right-sided occipito-temporal channels appears (590 to 635 ms, p = 0.0040). However, these evoked responses did not differ significantly between reversals (at p<0.05; BF_10_ < 0.79 for each ROI at -50, 0, 50 ms; Figure 4B) suggesting that the dynamics of the evoked responses is independent of the precise perceptual interpretation.

### Time-frequency power differences between reversals

We quantified rhythmic EEG signals using time-frequency (TF) analysis. This revealed a systematic decrease in alpha (8-12 Hz) and beta (13-30 Hz) power around the reversal (Figure 5A). To facilitate testing for a difference between reversals we grouped the different electrodes into five regions of interest (ROI; topography in Figure 5) and contrasted the two reversals within each ROI (Figure 5B-F). This revealed significant differences in alpha and beta power over occipital (9-20 Hz, at 0.22 to 0.82 s, t = -4.173, p = 0.0018; 13-19 Hz, 0.82 to 0.96 s, t= -4.159, p = 0.0488), parietal (9-25 Hz, 0.04 to 0.7 s, t = -4.8507, p = 0.0010; 12-21 Hz, 1.66 to 2.0 s, t = -4.3782, p = 0.0126), central (9-14 Hz, 0.32 to 0.56 ds, t = -3.7282, p = 0.0432; 15-21 Hz, 1.74 to 2.0 s, t = -5.7101, p = 0.0074), frontal (10-15 Hz, 0.18 to 0.78 s, t = -3.9212, p = 0.0036), and temporal electrodes (10-21 Hz, 0.16 to 0.5 s, t = -4.5511, p = 0.0142; 12-25 Hz, 1.7 to 1.98 s, t = -5.4548, p = 0.0060). Alpha and beta power were higher for the to-non-dominant reversal. We also found differences in theta power prior to the reversal. Theta power was higher for the to-non-dominant reversal over occipital (4 Hz, -0.78 to -0.04 s, t = - 4.440, p = 0.030), parietal (4-5 Hz, -0.82 to -0.04 s, t = -5.339, p = 0.0160), and frontal (4 Hz, - 0.92 to -0.14 s, t = -4.9562, p = 0.0208) electrodes. Clusters encompassing frequency ranges from beta to low gamma prior to the reversal report occurred over parietal (19-34 Hz, -0.72 to -0.52 s, t = -4.4362, p = 0.0246), central (15-29 Hz, -0.6 to -0.44 s, t = -4.4928, p = 0.0114), and temporal (20-34 Hz, -1.1 to -0.94 s, t = -4.6743, p = 0.0216; 20-40 Hz, -0.62 to -0.5 s, t = - 4.6684, p = 0.0220) electrodes, and also here power was higher for the to-non-dominant reversal.

### Heartbeat evoked potentials do not differ between reversals

The heartbeat evoked potential averaged across all heartbeats revealed a fronto-parietal topography known from previous studies (Gray et al., 2007; Cambi et al., 2021; review by Park & Blanke, 2019); though some studies also report a reversed polarity (Montoya et al., 1993; Pollatos & Schandry, 2004; Gautier et al., 2025). This comprises an initial positivity over posterior regions that was not significant (no cluster; p > 0.05) and a prolonged frontal negativity (cluster of the entire time range; p = 0.0002). Contrasting the HEP between the two reversals revealed no significant differences (at p<0.05; Figure 6B). Bayes factors calculated at different time points were mostly below 1 (except at 320 ms over parietal channels, BF_10_ =2.18), but none provided evidence in favor of a difference.

**Figure 6:**
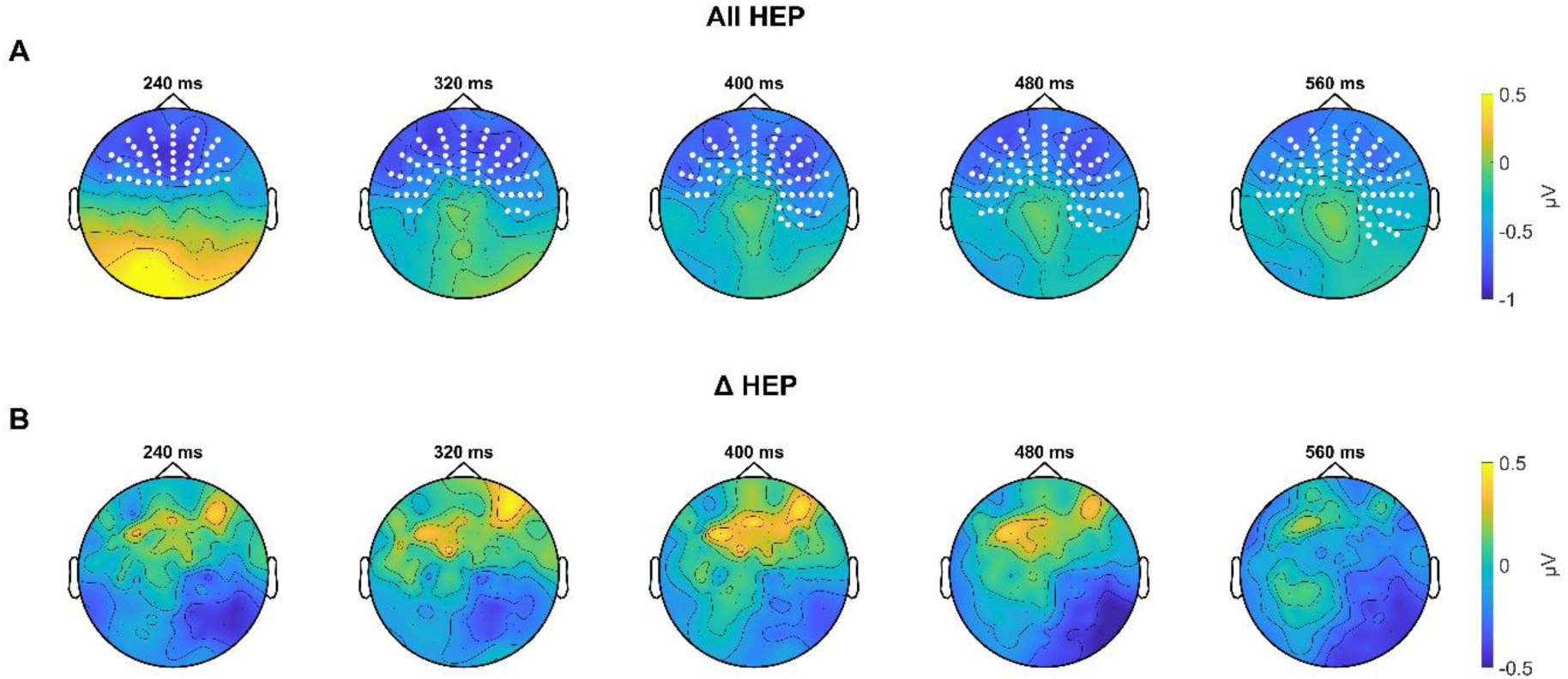
**A)** Average HEP across all heartbeats. Topographies show the average of a ± 40 ms window centered on the time relative to R peak indicated above each topography. The signal varied significantly from zero over fronto-temporal channels indicated with white dots (cluster p = 0.0002). **B)** Difference between HEP of the last full heartbeat preceding reversals (to-dominant minus to-non-dominant). Cluster permutation revealed no significant differences.

## Discussion

We investigated whether and how peripheral and central physiological signals exhibit systematic changes around spontaneous changes in perception of the Necker cube. In our data, respiration rate and heart rate changed systematically around reversals but did not differ between the two types of reversals. The same applied to EEG evoked responses as well as heartbeat evoked potentials. In contrast, the phase alignment of respiration, pupil size as well as the power of rhythmic EEG activity systematically differed between the two reversals. This suggests that signals relating to the body’s and brain’s momentary state are predictive of the perceptual interpretation of ambiguous stimuli and relate to the processes implementing the endogenously driven change in perception.

### Respiration and heart rate capture the temporal dynamics of reversals

The heart rate exhibited a biphasic pattern around reversals, decelerating prior to this and accelerating afterwards. This is reminiscent of the known phenomenon of anticipatory cardiac deceleration (Jennings & Woods, 1977). Since the spontaneous change in the interpretation of the Necker cube does not follow an explicit external cue, this deceleration must be driven by some intrinsic mechanism. In principle one could envisage respiratory sinus arrhythmia (RSA) as a cause, the systematic decrease in heart rate during expiration (Berntson et al., 1993). However, this seems unlikely given that respiratory phase locking prevailed only after the reversal, speaking against a clear relation between respiration phase and heat rate prior to the reversals. Further, the average respiratory phase after a reversal was mid to late expiration, which would imply a slowing in heart rate via the RSA, the opposite of what we observed. Some studies advocate that anticipatory cardiac deceleration comprises both a respiration dependent and independent component, arguing for a potential top-down regulation of vagal tone to induce cardiac slowing (Kingir et al., 2026). However, spontaneous reversals lack the external cue that is considered necessary for this phenomenon (Skora et al., 2022). One may also speculate that the previous perceptual reversal serves as an internal temporal reference inducing a cardiac deceleration prior to the subsequent reversal. However, this seems in conflict with the acceleration in heart rate seen after a reversal. In any case, the present data reveal specific dynamics of cardiac activity around spontaneous perceptual reversals that seems to be independent of the perceptual interpretation of the visual object.

Also, respiration frequency (or the duration of respiratory cycles) was significantly modulated around reversals. In our data respiration frequency systematically increased directly around reversals. Based on the time course and the slow nature of respiration one cannot say whether this is related to an upcoming change in perceptual interpretation or the fact that participants reported, via motor action, the perceived change in perception. However, both an increase in heart rate and an increase in respiration frequency reflect a level of increased metabolic activity that could be related to an increase in arousal, or alertness (Peters et al., 1998; Grassman et al., 2016; Charles & Nixon, 2019). Such a change in arousal around perceptual reversals is also reflected in the modulation of alpha band EEG activity, and pupil size (see below). Interestingly, neither heart rate nor respiration frequency differed between the two types of reversals, suggesting that the modulation of these signals is related to endogenous changes in perception or the reporting of this but not the specific perceptual interpretation of the Necker cube.

### Phase locking of respiration is sensitive to the reversal

Recent work on respiration shows that respiration phase tends to align to the time course of externally presented stimuli (Kluger et al., 2021; Della Penna et al., 2026; Chalas et al., 2026; Kayser et al., 2026; Harting et al., 2025; Stetza et al., 2025). As this phase locking can appear already prior to expected stimuli (Kluger et al., 2021; Stetza et al., 2025; Harting et al., 2025; Kayser et al., 2026), this may reflect some form of active sensing, triggered by the predictability of the external stimuli (Kluger et al., 2021; Della Penna et al., 2026; Chalas et al., 2026). The spontaneous changes in perception studied here lack this temporal predictability, since the reversal is not triggered externally and the distribution of perceptual stabilities has large variability. Not surprisingly, the respiratory phase-locking in the present data only followed the reversal. One possible cause of this could be related to the motor action of reporting the reversal. Respiratory alignment to motor actions has been shown in different experimental conditions, both self-initiated stimulus and externally paced (Perl et al., 2019; Park et al., 2020; Park et al., 2022; Shibata & Ohira, 2026; Raßler et al., 1996; Li & Laskin, 2006). Additionally, one could have expected a stronger phase locking for the to-dominant reversal, since this on average occurs on the same time scale (around 4.10 s; c.f. Fig 2) as the average duration of respiration cycles (which was 4.06 s). The match of these time scales could in principle facilitate respiratory alignment. However, while respiration frequency did not differ between the two reversals, phase locking was stronger for the reversal to the non-dominant interpretation, speaking against this hypothesis. An alternative interpretation is that the non-dominant interpretation is associated with higher sensory or decision uncertainty, and this triggers the alignment of respiratory resources to cope with this uncertainty, in line with respiration serving as a tool to allocate cognitive resources for sensory-motor challenges (Kayser et al., 2026).

### Pupil size modulates differentially around reversals

In line with previous studies, we found that pupil size was strongly modulated around reversals, with the pupil constricting prior to a reversal and dilating subsequently (Einhäuser et al.,2008; Hupé et al., 2009; Brascamp et al., 2021; Nakano et al., 2021). Since pupil size is linked to the activity of the autonomic nervous system (Aston-Jones & Cohen, 2005; Laeng et al., 2012; Viglione et al., 2023), this can be interpreted as a modulation of arousal around the reversal. However, previous studies have also speculated that the motor action of reporting the reversal may contribute to the subsequent dilation (Hupé et al., 2009; Brascamp et al., 2021).

Noteworthy, pupil size also differed between the two reversals and was modulated more strongly after the to-dominant reversal featuring a stronger dilation following this. Phasic changes in pupil size have been associated with a number of factors, including perceptual surprise, the commitment to a specific decision, or the engagement of top-down cognitive effort. For example, experiencing infrequent (rare) stimuli was shown to evoke larger pupil dilations than high-frequency (standard) stimuli (Murphy et al., 2014; Mathôt, 2018). Based on this one could have expected a stronger pupil dilation following the to-non-dominant reversal. Similarly, pupil size has also been associated with prediction errors, with larger pupil dilation following larger prediction errors (Preuschoff et al., 2011; Koenig et al., 2018; Harris et al., 2022). Again, the to-non-dominant reversal should induce a larger prediction error and larger pupil size, but this is not what we observed. To explain a larger pupil dilation following the to-dominant reversal, one could speculate that processes related to decision certainty, commitment and conflict resolution drive this, since pupil dilation has been also associated with decision confidence and meta-cognitive processes (De Gee et al., 2014; Lempert et al., 2015). The return to the dominant interpretation may be associated with higher decision confidence, and this reflects in pupil size. This would also be in line with the speculation that stronger respiration phase locking for the non-dominant interpretation is associated with greater uncertainty.

### Oscillatory dynamics but not evoked responses are reversal-specific

The time-locked event-related potentials revealed a significant modulation around reversals. This not only included central activity at or just following the button press that could reflect motor processes, but also fronto-occipital differences around 300 ms prior to the indicated reversal as well as components up to 600 ms following this. This points to a number of physiological processes that underlie and accompany the reversal. One could speculate that the frontal activity 300 ms prior to the reversal report may be linked to the destabilization of the current percept, the resolution of competing perceptual interpretations, or the preparation of the subsequent perceptual report via a motor action (İşoğlu-Alkaç et al., 1998; İşoğlu-Alkaç et al., 2000; Wilson et al., 2023). Similarly, the late activity occurring at 650 ms could be related to the stabilization of the new percept. However, in contrast to studies relying on paradigms with flashed stimuli or intermittent tasks designs (Kornmeier & Bach, 2006; Pitts et al., 2009; Kornmeier & Bach, 2012), the interpretation of these ERP markers is more difficult and possibly also affected by higher inter- and intraindividual variability in the time between of becoming aware of the change in perceptual interpretation and subsequently reporting this change. Furthermore, in our data these evoked responses did not differ between the two reversals, suggesting that these processes are not specific to the momentary perceptual interpretation of the cube but mainly the process of switching and indicating the percept.

A similar observation held for the heartbeat evoked potentials. The time course of the HEP activity comprised strong occipital and central components, similar to previous reports (Gray et al., 2007; Cambi et al., 2021; review by Park & Blanke, 2019), although we note that the specific pattern of HEPs also seems to differ across studies (Montoya et al., 1993; Pollatos & Schandry, 2004; Gautier et al., 2025). The HEP did not differ significantly between reversals, suggesting that this marker of interoceptive cardiac processing is not affected by the perceptual interpretation of the cube. Also, since we could include only the data from 23 participants in this analysis, the results on the HEP have less statistical power compared to the other data reported here.

The analysis of rhythmic EEG activity revealed a number of significant effects around the reversals. Similar to the reports of previous studies (İşoğlu-Alkaç & Strüber, 2006; Piantoni et al., 2010; Yokota et al., 2014; Piantoni et al., 2017; Drew et al., 2022; Mokri et al., 2025; Drew et al., 2026), we observed a general decrease in alpha and beta power prior to the reversal and subsequent increase (Figure 5A). Prominently, alpha and beta band activity differed between reversals, in line with previous studies linking these time scales of neural activity to endogenous changes in perception (Piantoni et al., 2010; Yokota et al., 2014; Piantoni et al., 2017; Drew et al., 2022; Mokri et al., 2025; Drew et al., 2026). These differences in alpha and beta power emerged only after the reversal report, lasting several 100 ms and occurred in all ROIs. Both alpha and beta power were higher for the to-non-dominant reversal. Alpha activity has been associated with arousal and cortical excitability, with higher arousal being associated with reduced alpha power (Romei et al., 2008; Romei et al., 2010; Jensen & Mazaheri, 2010; Iemi et al., 2017). The transition from a stable neural representation to a dynamic one hence seems to be facilitated by a modulation of neural excitability and inhibitory processes (İşoğlu-Alkaç & Strüber, 2006; Piantoni et al., 2010; Piantoni et al., 2017; Drew et al., 2022; Mokri et al., 2025; Drew et al., 2026). Importantly, in this line of thought our data suggest that the to-dominant reversal is associated with higher arousal or excitability, which directly fits with the results on pupil size, which was more dilated for this type of reversal. As discussed above, however also other factors than arousal could explain this difference in alpha power, such as differences in decision certainty or other meta-cognitive processes (De Gee et al., 2014; Lempert et al., 2015), whereby the transition to the dominant interpretation may reflect the return to a high-confidence internal state.

Beta band activity has been associated with maintaining the status quo (Engel & Fries, 2010) but also the top-down endogenous reactivation of specific mental representations (Spitzer & Haegens, 2017). Since the non-dominant interpretation is possibly less supported by automatic bottom-up process, increased top-down feedback to sensory cortices is required to establish and sustain this non-dominant interpretation. This is also in line with work on predictive coding, where beta band activity establishes descending feedback (Bastos et al., 2012; Bastos et al., 2015). Beta band activity also emerges in relation to motor preparation and execution. While the beta band effects did not emerge over central electrodes, we cannot exclude a contribution of motor processes to these effects. However, since the motor action of reporting a reversal was very similar for both interpretations of the cube, a pure motor related effect seems very unlikely and processes related to top-down feedback are the more parsimonious explanation. Finally, those differences in alpha / beta power emerging very late in our analysis window (1.66 to 2.0 s) could possibly be related to processes already pertaining to the subsequent reversal.

We also observed a difference in theta power between reversals, which emerged over frontal, parietal, and occipital electrodes prior to the reversal, similar as theta-band effects reported in previous studies in binocular rivalry (Drew et al., 2022; Drew et al., 2026). The differences in theta band power started around one second prior to the reversal, with the earliest cluster located over frontal electrodes (starting -0.92 s), followed by the parietal cluster (starting - 0.82 s) and then the occipital cluster (starting -0.72 s). Theta band activity has been linked to conflict monitoring and conflict resolution during information processing (Cohen, 2014) and in our data theta power was higher for the to-non-dominant reversal. Hence this could be interpreted as a stronger engagement of conflict resolution when seeing the dominant cube while being in the process of switching to the non-dominant interpretation. The reversal towards the non-dominant interpretation may simply be associated with stronger needs to resolve conflicts than the reversal to the predominant interpretation.

Finally, we also found differences in gamma power between reversals. These also emerged prior to the reversal report and over multiple sites. Some previous studies have linked gamma activity to upcoming spontaneous switches in perception, e.g. in the context of binocular rivalry (Doesburg et al., 2005; Doesburg et al., 2009; Ehm et al., 2011), suggesting that the underlying processes may be related to a change in sensory encoding. Indeed, gamma power was higher for the to-non-dominant reversal. In the context of predictive coding, gamma band activity has been associated with feed-forward processing (Bastos et al., 2012; Bastos et al., 2015), suggesting that the neurophysiological activity prior to a change to the non-dominant interpretation is associated with higher feed-forward processing, possibly to establish this interpretation against the default interpretation. This would be in line with the reversal to the non-dominant interpretation also requiring the resolution of conflicts between bottom-up sensory input and top-down expectations, which is reflected in theta activity. While the functional and mechanistic interpretations of these changes in oscillatory dynamics certainly remain speculative and post-hoc, the data nevertheless reveal the richness of neurophysiological processes that differentiate the two interpretations of the Necker cube.

## Conclusion

Taken together, our findings suggest that spontaneous perceptual reversals are accompanied by coordinated changes across peripheral and cortical signals. Rather than reflecting independent physiological processes, these signals may represent different manifestations of a common change in neural state associated with the need to resolve conflicts between two potential interpretations of the same object and to establish a new perpetual interpretation. While the effects observed in the different signals differ in their timing, the present data cannot dissociate potential causal relationships between these, nor the precise underlying mechanisms. The timing and magnitude of reversal differences in these signals may in part reflect the different intrinsic time scales of the respective physiological systems. This may in part explain why cardiac, respiratory, pupillary, and neural measures exhibit distinct temporal relationships to spontaneous perceptual reversals. All in all our data highlight the value and need to simultaneously assess multiple peripheral and cortical signals in relation to perception and cognition, as these provide complementary insights into how changes in cortical state are reflected across interacting physiological systems.

## Acknowledgement

We would like to thank Sepideh Mirzaei and Jessalyn Dvorak for their help in data collection.

## Author contributions

L.S. and C.K. designed research; L.S. performed research; C.K. and L.H. contributed unpublished reagents/analytic tools; L.S. and C.K. analyzed data; L.S. and C.K. wrote the paper.

## Data availability

Preprocessed data used to generate figures and to perform statistics are available from the corresponding author upon request.

## AI usage

OpenAI has been used for language and grammar improvements. AI tools were used to assist with structuring and streamlining the analysis code, including improving code organization and reducing redundancy. All analyses and methodological decisions were performed and verified by the authors.

